# Serum levels and tumour expression of leptin and leptin receptor as promising clinical biomarkers of specific feline mammary carcinoma subtypes

**DOI:** 10.1101/2020.04.27.064600

**Authors:** Andreia Gameiro, Catarina Nascimento, Ana Catarina Urbano, Jorge Correia, Fernando Ferreira

**Author notes:** Correspondence (F.F.); / Ext.: 431234 (F.F.).

## Abstract

Obesity is a risk factor for human breast cancer, being associated with increased serum levels of leptin. In cat, although obesity is a common nutritional disorder, the role of leptin and its receptor in mammary carcinoma is unknown. In this study, serum levels of leptin and leptin receptor (ObR) were evaluated in 58 cats with mammary carcinoma and compared with healthy controls by ELISA, as tumour expression by immunohistochemistry. Results showed that the free leptin index is decreased in cats with mammary carcinoma (p=0.0006), particularly for those with luminal B and HER2-positive disease that showed significantly lower serum leptin levels (p<0.0001 and p<0.005, respectively). Serum leptin levels above 4.17 pg/mL were associated with ulcerating tumours (p=0.0005) and shorter DFS (p=0.0217). Elevated serum ObR levels were found in all cats with mammary carcinoma (p<0.0001), with levels above 16.89 ng/mL being associated with smaller tumours (p=0.0118), ER-negative status in HER2-positive tumours (p=0.0291) and increased serum levels of CTLA-4 (p=0.0056), TNF-α (p=0.0025), PD-1 (p=0.0023) and PD-L1 (p=0.0002). In tumour samples, leptin was overexpressed in luminal B and triple-negative carcinomas (p=0.0046), whereas ObR was found overexpressed in luminal B samples (p=0.0425). Altogether, our results reinforce the importance of feline mammary carcinoma in comparative oncology.

## 1. Introduction

The feline mammary carcinoma (FMC) is a high prevalence disease (12 to 40% of all tumours in cat) that shows similar clinicopathological features to human breast cancer [1], supporting its use in comparative oncology studies [2] [3]. Likewise, obesity is a common nutritional disorder in cat, with higher prevalence in indoor and sterilized animals above three years of age [4]. In humans, obesity induces a chronic inflammatory status, being a risk factor for breast cancer [5] [6] [7].

Leptin is a 16 kDa adipocytokine, encoded by the *obese* gene and involved in the central regulation of food intake, energy homeostasis, modulation of reproductive function and peripheral metabolic processes, such as breast/mammary gland development, cellular proliferation and angiogenesis. In tissues and serum, leptin expression is modulated by fat mass, with healthy cats showing lower serum leptin levels than obese animals [4], as reported in humans [5] [6]. Interestingly, although this protein is mainly secreted by adipocytes, it can be also expressed by pathologically altered cells, such as cancer cells [8] [9]. Thus, malignant cells can regulate their metabolic activities [10], promoting uncontrolled cell growth, migration, invasion and angiogenesis [11] [7], and downregulating apoptosis through a Bcl-2-dependent mechanism [10] [12]. Accordingly, leptin overexpression is detected in breast cancer cells and neighbouring adipocytes, contrasting with normal breast glandular epithelial cells [13] [7], suggesting an oncogenic role for this adipocytokine [6]. Furthermore, studies in human breast cancer patients showed that leptin overexpression has paracrine effects, not always reflected in serum levels, although associated with more aggressive tumours and therapy resistance [13]. Additionally, in overweight human patients a positive correlation was found between leptin overexpression in the tumour microenvironment and oestrogen receptor (ER) positive breast cancer, and with a human epidermal growth factor receptor 2 (HER 2)-positive status frequently related to a more invasive tumour phenotype [14].

In parallel, the leptin receptor (ObR, 150-190 kDa) was found to be involved in innate and adaptive immunity [15], being expressed in several organs, including breast and peripheral tissues, as well as in adipocytes [16] [17] and immune cells. ObR is constituted by an extracellular N-terminus domain, a transmembrane domain, and a cytoplasmic C-terminus domain. Upon leptin ligation, ObR homodimerizes and the associated JAK monomer is auto phosphorylated to activate the downstream signalling pathways [8]. The soluble ObR form is a 146 kDa protein [18] that could be generated by cellular apoptosis or by the proteolytic cleavage of the extracellular anchored protein domain, with this shedding being more frequent in shorter intracellular isoforms. In serum, the ObR modulates the leptin bioavailability, being decreased in obese humans [8]. In breast cancer patients, ObR is overexpressed independently of the ER status [6], being correlated with low overall survival (OS) [9]. Furthermore, the ratio between leptin/ObR serum levels (free leptin index – FLI) is considered an useful predictor of leptin activity, reflecting the individual metabolic status [19] and when increased, is an important risk factor for breast cancer development [20]. In parallel, studies in breast cancer patients found an association between leptin and ObR overexpression with a chronic inflammatory status, conditioning T-cell immune responses (increase Th1- and decrease Th2-responses) [21] and the activation of immune checkpoint inhibitors [16]. Indeed, some studies in humans have shown a positive correlation between overexpression of leptin and ObR with several immunomodulatory molecules (e.g. Cytotoxic T-Lymphocyte Associated Protein 4 - CTLA-4; Tumor Necrosis Factor α - TNF-α; Programmed Cell Death-1 - PD-1 and Programmed Cell Death-ligand 1 - PD-L1) [22] [23]. While the CTLA-4 is a protein related to the inflammatory response that is increased in breast cancer patients, contributing for the immune response downregulation [24], the TNF-α is a pro-inflammatory cytokine that induces apoptosis promoted by the absence of leptin [25]. Moreover, the overexpression of PD-1 in T-cells is associated with ObR overexpression in humans with distinct tumour types [26], induced through the AKT pathway activation by oestrogens [27] and being responsible for the PD-1 mediated T-cell dysfunction [28].

As mentioned above, obesity is associated with increased leptin levels, which induces resistance to chemotherapy [29] [30]. Therefore, the leptin/ObR axis has been widely studied [31] as a target for a adjuvant therapy, not only in ER-positive tumour status [29], but also in triple-negative tumours [32], in which the lack of hormonal receptors reduces the therapeutic options. The use of leptin antagonists, allows the blocking of the leptin receptor, leading to a downregulation of the leptin downstream pathways, some of them known as oncogenic (e.g. Wnt and STAT3) [29] [32] [31].

To the best of our knowledge, this study is the first to evaluate the leptin and ObR serum levels and tissue expression in cats with mammary carcinoma. The main goals of this study were as follows: 1) to compare the serum leptin and ObR levels between cats with mammary carcinoma stratified by molecular subtype and healthy controls, using ELISA; 2) to investigate the leptin and ObR expression in mammary carcinoma samples and compare with normal mammary tissues, using immunohistochemistry (IHC); 3) to search for associations between serum leptin/ObR levels and leptin/ObR IHC scores in tumour mammary tissues; 4) to test for associations between serum leptin/ObR levels and several clinicopathological features, in order to evaluate the utility of leptin and ObR as diagnostic and/or prognosis biomarkers or as promising drug targets for feline mammary carcinoma.

## 2. Materials and Methods

### 2.1. Animal population

Tumour tissue and serum samples were collected from 58 female cats with fully documented history of FMC that underwent mastectomy and 24 serum samples from healthy cats presented for elective ovariohysterectomy, at the Teaching Hospital of the Faculty of Veterinary Medicine, University of Lisbon. The animals were anesthetized before surgical procedures and blood samples were collected with no interference in animal well-being, with the procedures involving manipulation of animals being consented by the owners. Briefly, all tissue samples were embedded in paraffin after fixation in 10% buffered neutralized formalin (pH 7.2), during 24-48 hours, while serum samples were separated from clotted blood by centrifugation (1500g, 10min, 4°C) and stored at −80°C until further use. All samples that showed haemolysis were discarded, as recommended [1] [33].

For each animal enrolled in the study, the clinicopathological data were recorded, including age, breed, body weight, reproductive status and contraceptive administration, treatment (none, mastectomy or mastectomy plus chemotherapy), number, location and size of tumoral lesions, histopathological classification (ER status, PR status, HER2 status and Ki-67 index), malignancy grade, presence of tumour necrosis, lymphatic invasion, lymphocytic infiltration, cutaneous ulceration, regional lymph node involvement and clinical stage (TNM system) (Table 1). Regarding the molecular subtyping of feline mammary carcinomas [34], animals were stratified in luminal A (n=10), luminal B (n=17), HER2-positive (n=15) and triple-negative (n=16) groups.

**Table 1.**
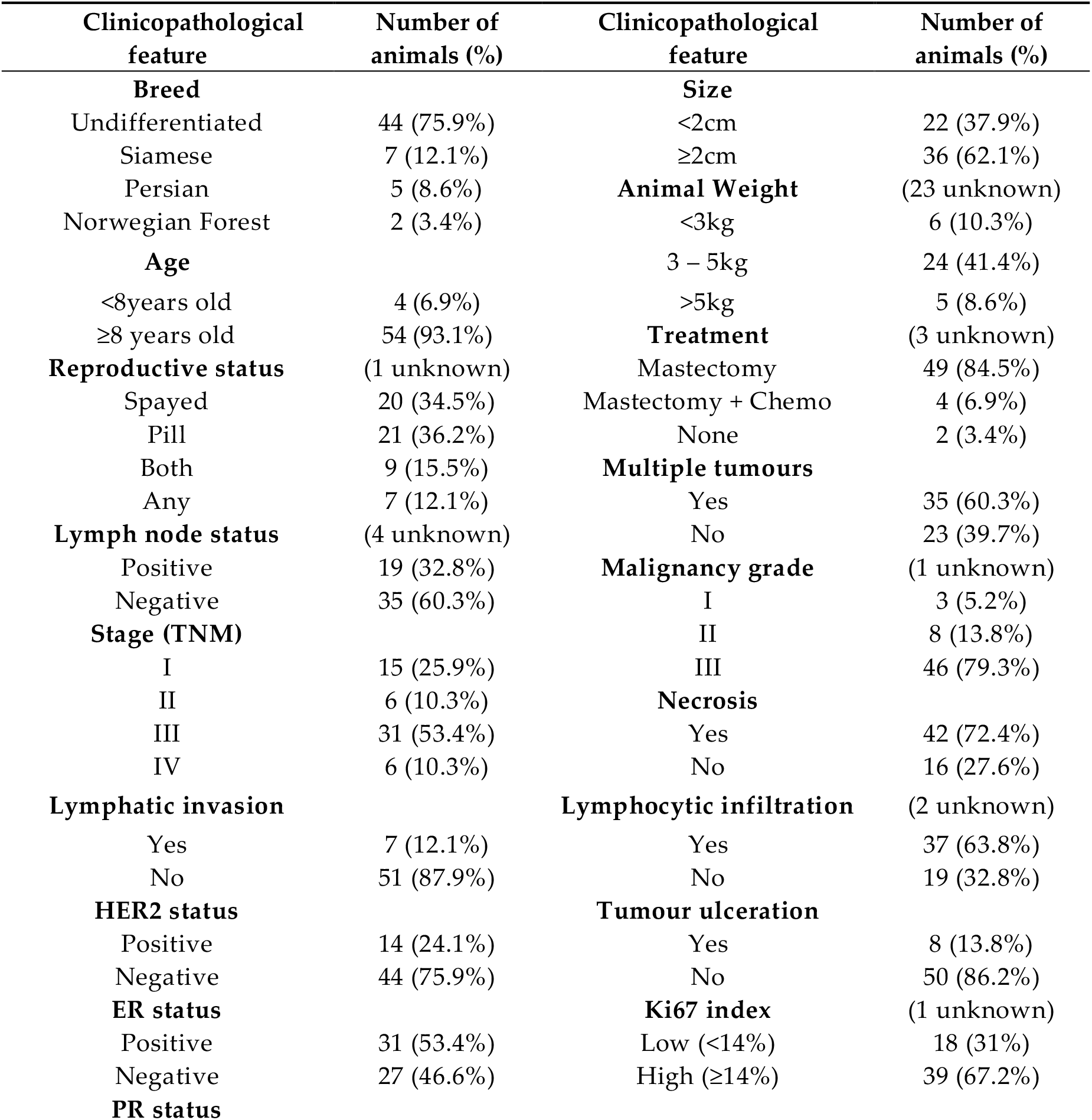
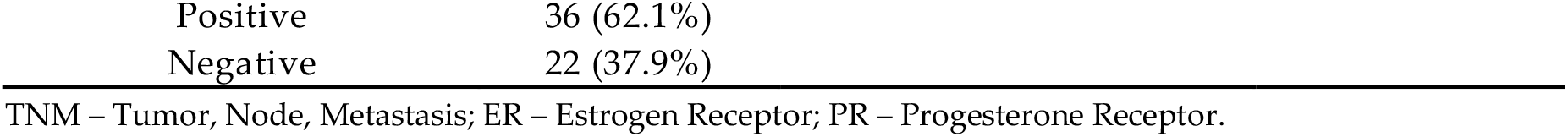
Clinicopathological features of female cats with mammary carcinomas enrolled in this study (n=58).

| Clinicopathological feature | Number of animals (%) | Clinicopathological feature | Number of animals (%) |
| --- | --- | --- | --- |
| **Breed** |  | **Size** |  |
| Undifferentiated | 44 (75.9%) | <2cm | 22 (37.9%) |
| Siamese | 7 (12.1%) | ≥2cm | 36 (62.1%) |
| Persian | 5 (8.6%) | **Animal Weight** | (23 unknown) |
| Norwegian Forest | 2 (3.4%) | <3kg | 6 (10.3%) |
| **Age** |  | 3 – 5kg | 24 (41.4%) |
| <8years old | 4 (6.9%) | >5kg | 5 (8.6%) |
| ≥8 years old | 54 (93.1%) | **Treatment** | (3 unknown) |
| **Reproductive status** | (1 unknown) | Mastectomy | 49 (84.5%) |
| Spayed | 20 (34.5%) | Mastectomy + Chemo | 4 (6.9%) |
| Pill | 21 (36.2%) | None | 2 (3.4%) |
| Both | 9 (15.5%) | **Multiple tumours** |  |
| Any | 7 (12.1%) | Yes | 35 (60.3%) |
| **Lymph node status** | (4 unknown) | No | 23 (39.7%) |
| Positive | 19 (32.8%) | **Malignancy grade** | (1 unknown) |
| Negative | 35 (60.3%) | I | 3 (5.2%) |
| **Stage (TNM)** |  | II | 8 (13.8%) |
| I | 15 (25.9%) | III | 46 (79.3%) |
| II | 6 (10.3%) | **Necrosis** |  |
| III | 31 (53.4%) | Yes | 42 (72.4%) |
| IV | 6 (10.3%) | No | 16 (27.6%) |
| **Lymphatic invasion** |  | **Lymphocytic infiltration** | (2 unknown) |
| Yes | 7 (12.1%) | Yes | 37 (63.8%) |
| No | 51 (87.9%) | No | 19 (32.8%) |
| **HER2 status** |  | **Tumour ulceration** |  |
| Positive | 14 (24.1%) | Yes | 8 (13.8%) |
| Negative | 44 (75.9%) | No | 50 (86.2%) |
| **ER status** |  | **Ki67 index** | (1 unknown) |
| Positive | 31 (53.4%) | Low (<14%) | 18 (31%) |
| Negative | 27 (46.6%) | High (≥14%) | 39 (67.2%) |
| **PR status** |  |  |  |

|  |  |
| --- | --- |
| Positive | 36 (62.1%) |
| Negative | 22 (37.9%) |

TNM – Tumor, Node, Metastasis; ER – Estrogen Receptor; PR – Progesterone Receptor.

### 2.2. Measurement of serum leptin, ObR, CTLA-4, TNF-α, PD-1 and PD-L1 levels

The serum levels of leptin, ObR, CTLA-4, TNF-α, PD-1 and PD-L1 were measured by using commercial ELISA-based kits from R=D Systems (Minneapolis, USA; DY389, DY398-05, DY476, DY2586, DY1086, DY156, respectively) and following the manufacture’s recommendations. After collection, the serum samples were kept stored at −80°C and thawed shortly before use. For each assay, a standard curve was generated by using serial dilutions of the recombinant proteins from kits. For leptin, ObR, PD-1 and PD-L1 the r^2^ values were calculated using a quadratic regression (r^2^= 0.9976, for leptin, r^2^= 0.9632, for ObR, r^2^=0.99 for PD-1 and r^2^=0.96 for PD-L1), whereas serum CTLA-4 and TNF-α concentrations were determined by using a curve-fitting equation (r^2^>0.99), as previously reported [35].

Briefly, a 96-well plate was prepared by adding the capture antibody to each well and incubate overnight. Plates were then treated with 1% bovine serum albumin (BSA) in phosphate buffered saline (PBS), for 1 hour, to prevent nonspecific binding. Standards and diluted serum samples were added to sample wells and incubated for 2 hours at room temperature (RT), before the incubation of the detection antibody for 2 hours at RT. Afterwards, the streptavidin-conjugated to horseradish peroxidase (HRP) was added to each well and incubated at RT, for 20 minutes, before the addition of the substrate solution in 1:1 H_2_O_2_ and tetramethyl-benzidine to each well (20 minutes, at RT, in the dark). The reaction was interrupted by adding a stop solution (2NH_2_SO_4_) and the absorbance was measured by a spectrophotometer (FLUOStar OPTIMA, Microplate Reader, BMG, Ortenberg, Germany), using 450 nm as the primary wavelength and 570 nm as a reference wavelength.

### 2.3. Assessment of the leptin and ObR status by immunohistochemistry (IHC)

Initially, the feline mammary carcinoma formalin fixed paraffin-embedded (FFPE) samples were stained with haematoxylin-eosin to select a representative tumour area and a normal tissue area to be used as control (n=20). FFPE samples were sectioned in slices with 3 μm thickness (Microtome Leica RM135, Newcastle, UK) and mounted on a glass slide (SuperFrost Plus, Thermo Fisher Scientific, Massachusetts, USA). On PT-Link module (DAKO, Agilent, Santa Clara, USA), samples were deparaffinized, hydrated and antigen retrieval was performed, for 20 minutes, at 96°C, using Tris-EDTA buffer, pH 9.0 (EnVision™ Flex Target Retrieval Solution High pH, Dako). Then, slides were cooled for 30 minutes at RT and immersed twice, for 5 minutes in distilled water. IHC technique was performed with commercial solutions from the Novolink™ Max Polymer Detection System Kit (Leica Biosystems, Newcastle UK). Before antibody incubation, tissue samples were treated to block the endogenous peroxidase activity for 15 minutes, and the unspecific antigenic recognition was inhibited for 10 minutes. Finally, tissue samples were incubated at RT for 1 hour, in a humidified chamber, with the following primary antibodies: anti-leptin antibody (ab3583, AbCAM, Cambridge, UK) and anti-ObR antibody (ab104403, AbCAM), both diluted at 1:200. The slides were washed twice, for 5 minutes, between all the incubation steps, using a PBS solution at pH 7.4. Then, the detection polymer was incubated for 30 minutes, at RT, and detection was performed using diaminobenzidine (DAB substrate buffer and DAB Chromogen, Leica Biosystems) for 5 minutes. Later, samples were counterstained with Gills haematoxylin (Merck, New Jersey, USA) for 5 minutes, dehydrated in an ethanol gradient and xylene, and mounted using Entellan mounting medium (Merck).

To assess leptin and ObR immunoreactivity were used a scoring system previously reported [6] [13] [36] and the H-Score published by the American Society of Clinical Oncology (ASCO). The final IHC score was obtained by multiplying the positive cells (0=absence of staining; 1=all cells stained), by the highest staining intensity (Table 2), varying from 0 to 3, with tissue samples scored as 0 considered negative, while samples scored as 3 designated highly reactive. All slides were subjected to blind scoring, by two independent and experienced pathologists.

**Table 2.** Scoring criteria of immunostaining assay for leptin and ObR. Three microscopic fields were analyzed at 400x magnification.

| Stained tumour cells<br>(0-1) | Staining intensity |  |
| --- | --- | --- |
|  | Score | Interpretation |
|  | 0 | No staining |
|  | 1 | Weak |
|  | 2 | Moderate |
|  | 3 | Strong* |
| **Final IHC score = stained tumour cells x staining intensity score (0 – 3)** |  |  |

\* High protein reactivity

### 2.4. Statistical analysis

Statistical analysis was carried out using the GraphPad Prism software, version 5.04 (California, USA), with two-tailed p-values less than 0.05 considered statistically significant for a 95% confidence level (*p<0.05, **p<0.01 and ***p<0.001).

The non-parametric Kruskal-Wallis test was performed to compare leptin and ObR results between healthy cats and cats with mammary carcinoma stratified by tumour subtype. Receiver-operating characteristic (ROC) curves were performed to choose the optimal cut-off value for serum leptin and ObR levels, and to determine the specificity and sensitivity of the technique to diagnose the disease. The non-parametric Mann-Whitney test was used to compare the serum levels of both proteins with several clinicopathological features. Survival analysis was performed using the Kaplan-Meier test to evaluate the disease-free survival (DFS) in cats with mammary carcinomas. The correlations between serum ObR levels and serum concentrations of the inflammatory proteins CTLA-4, TNF-α, PD-1 and PD-L1 were investigated using the Spearman’s rank correlation coefficient.

## 3. Results

### 3.1. Cats with mammary carcinoma showed a reduced Free Leptin Index

The Free Leptin Index (FLI) was determined in the serum samples of cats with mammary carcinomas and compared with healthy animals. Results obtained showed that cats with disease had a significantly lower FLI than control group (0.44 vs. 0.86, p=0.0006, Figure 1).

**Figure 1.**
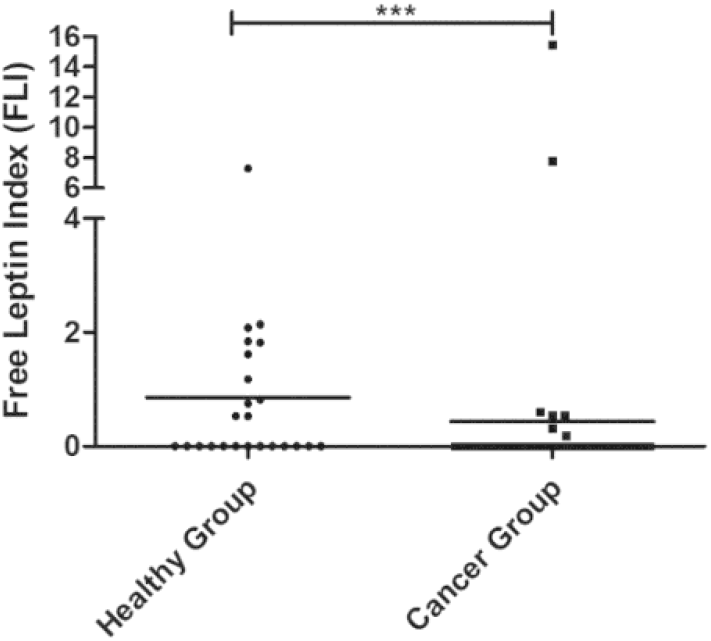
Dot plot diagram showing that the free leptin index (FLI) was significantly elevated in healthy animals that in cats with mammary carcinoma (p=0.0006).

In addition, results revealed that body weight did not influence serum leptin and ObR levels, both in the controls (p=0.0760 and p=0.8432, respectively, data not shown) and in the cancer group (p=0.3294 and p=0.9722, respectively, data not shown).

### 3.2. Cats with Luminal B or HER2-positive mammary carcinomas showed decreased serum leptin levels

Regarding serum leptin levels, results obtained showed that cats with luminal B or HER2-positive mammary carcinomas had lower serum leptin levels than healthy animals (0.00 pg/mL vs. 13.89 pg/ml, p<0.001; 0.83 pg/mL vs. 13.89 pg/mL, p<0.05, respectively, Figure 2A). The optimal cut-off value to predict mammary carcinoma was 4.17 pg/ml with an area under the ROC curve (AUC) of 0.7045±0.0757 (95% CI: 0.5561-0.8528, p=0.0103; sensitivity=96.9%; specificity=43.5%; Figure 2B). Further statistical analysis showed that elevated serum leptin levels were associated with tumour ulceration (p=0.0005, Figure 2C) and shorter DFS (117 vs. 314 days, p=0.0217, Figure 2D).

**Figure 2.**
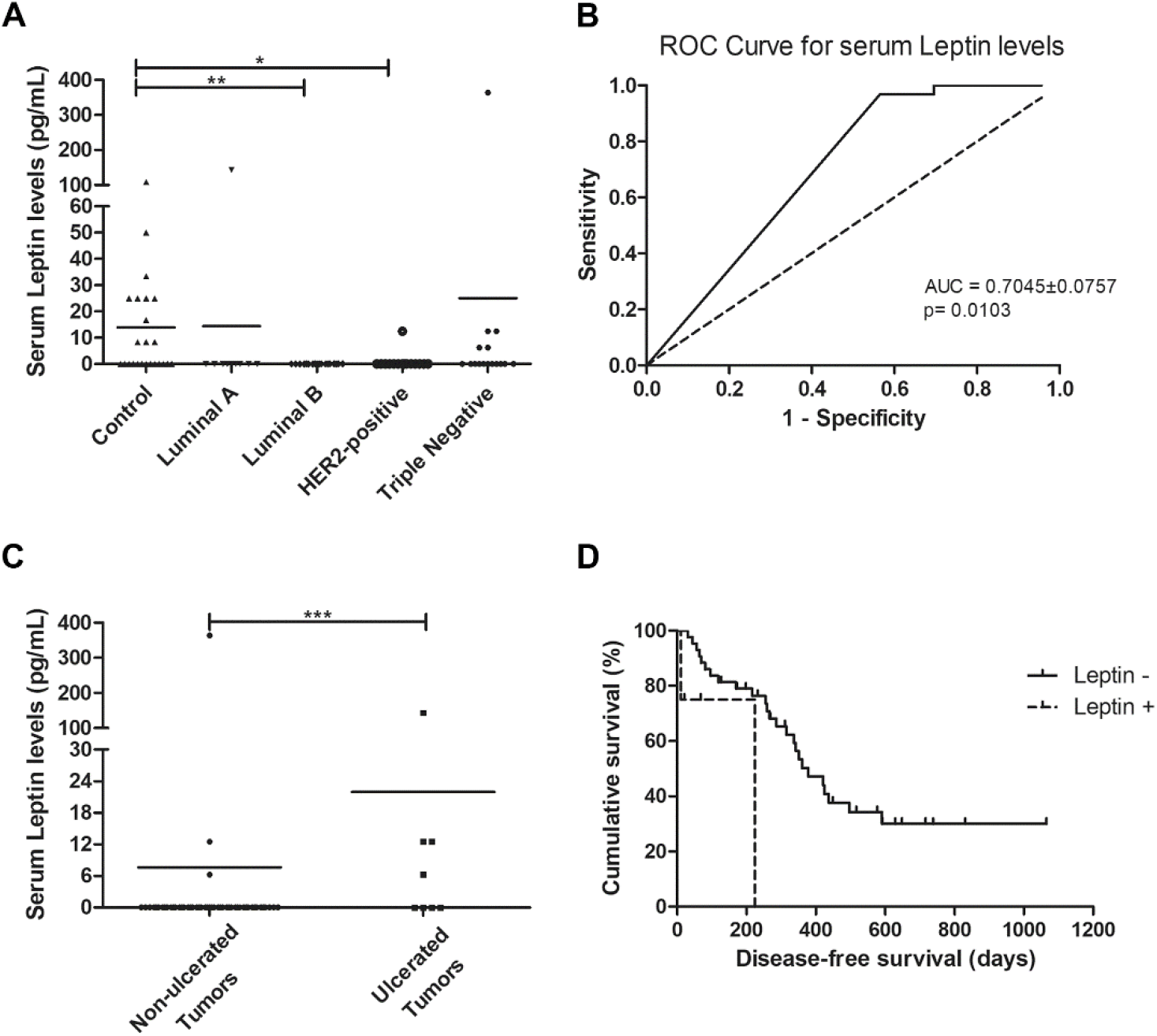
Cats with luminal B and HER2-positive mammary carcinomas showed decreased serum leptin levels, as cats had ulcerated tumours, with serum leptin levels above the cut-off level of 4.17 pg/mL being associated with shorter disease-free survival. **A)** Dot plot diagram showing the distribution of serum leptin levels (pg/mL) among healthy animals (control) and cats stratified by mammary carcinoma subtype. Significant decreased serum levels of leptin were found in cats presenting luminal B or HER2-positive subtypes in comparison to healthy animals (p=0.0025). **B)** The optimal cut-off of serum leptin levels to predict mammary carcinoma was determined to maximize the sum of the sensitivity and specificity (4.17 pg/mL; AUC=0.7045±0.0757, 95% CI: 0.5561-0.8528, p=0.0103; sensitivity=96.9%; specificity=43.5%). **C)** Dot plot diagram showing that serum leptin levels were significantly higher in cats with ulcerated tumours (p=0.0005). **D)** Cats with mammary carcinoma and serum leptin levels higher than 4.17 pg/mL had a lower DFS (p=0.0217). *p<0.05; **p<0.001; ***p<0.0001.

### 3.3. Cats with mammary carcinoma showed elevated serum levels of ObR and of mediators of inflammation

Considering the above results, the serum levels of the ObR were also evaluated. When the animals were grouped according to the tumour subtype, a significant difference was found between the mean ranks of at least one pair of groups (p<0.0001). Results revealed that serum ObR levels were significantly higher in animals with mammary carcinoma than in controls, independently of molecular subtype (control group 15.67 ng/ml; luminal A 23.04 ng/ml, p<0.0001; luminal B 20.18 ng/ml, p<0.001; HER2-positive 28.99 ng/ml, p<0.0001; triple-negative 21.70 ng/ml, p<0.0001; Figure 3A). Furthermore, the optimal cut-off value calculated for cats with mammary carcinoma was 16.89 ng/ml, with an AUC of 0.9408±0.0288 (95% CI: 0.8842-0.9973, p<0.0001; sensitivity=94.8%; specificity=87.0%; Figure 3B).

**Figure 3.**
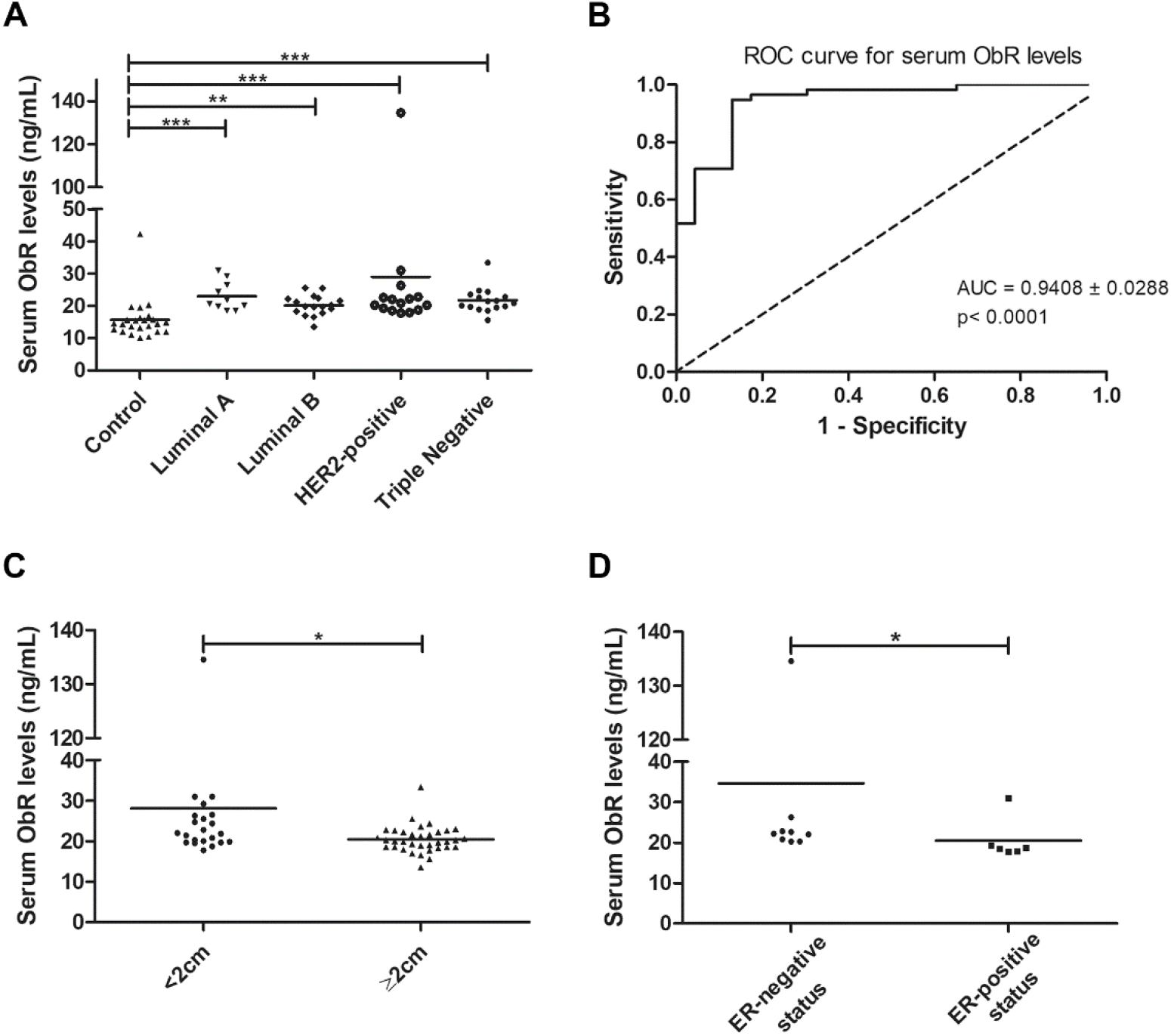
Cats with mammary carcinoma showed elevated serum ObR levels, with serum concentrations above 16.89 ng/mL being associated with smaller tumours and with HER-2 positive mammary carcinomas showing an ER-negative status. **A)** Dot plot diagram showing the distribution of serum ObR levels (ng/mL) in heathy animals (control) and in cats with mammary carcinoma stratified by molecular subtype. Significant higher serum levels of ObR were found in all tumour subtypes in comparison to healthy animals (p<0.0001). **B)** The optimal cut-off value of serum ObR levels to predict cats with mammary carcinoma was 16.89 ng/mL with an AUC of 0.9408±0.0288 (95% CI: 0.8842-0.9973, p<0.0001; sensitivity=94.8%; specificity=87.0%). **C)** Dot plot diagram showing that serum ObR concentrations were significantly low in tumours larger than 2 cm (p= 0.0118). **D)** Dot plot diagram for animals with HER2-positive mammary carcinoma displaying a positive association between higher serum ObR levels and ER-negative status (p=0.0291). *p<0.05; **p<0.001; ***p<0.0001.

In addition, elevated serum ObR levels were associated with smaller tumours (p=0.0118, Figure 3C) and with cats had HER2-positive mammary tumours showing an ER-negative status (p=0.0291, Figure 3D).

Moreover, the data obtained also showed a positive correlation between serum ObR levels and serum CTLA-4 (r=0.38, p=0.0056, Figure 4A), TNF-α (r=0.40, p=0.0025, Figure 4B), PD-1 (r=0.42, p=0.0023, Figure 4C) and PD-L1 (r=0.50, p=0.0002, Figure 4D) levels.

**Figure 4.**
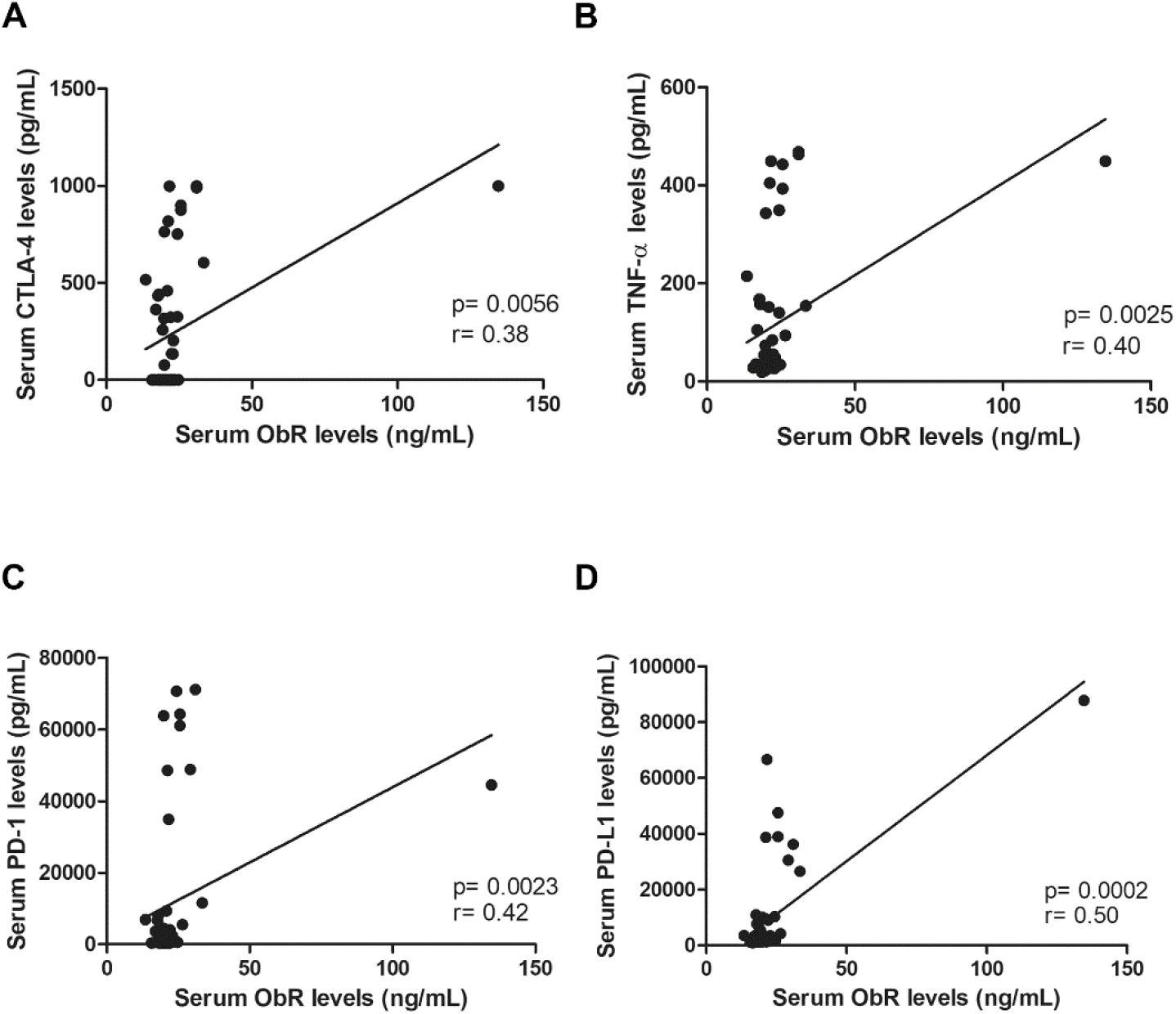
Serum ObR levels showed a positive correlation between serum ObR levels and **A)** serum CTLA-4 levels (p=0.0056), **B)** serum TNF-α levels (p=0.0025), **C)** serum PD-1 levels (p=0.0023) and **D)** serum PD-L1 levels (p=0.0002).

### 3.4. Leptin and ObR are overexpressed in Luminal B and triple-negative mammary carcinomas

The obtained results revealed that cats with luminal B or triple-negative mammary carcinoma showed a higher leptin IHC score than controls (1.93 vs. 1.34, p<0.05; 2.00 vs. 1.34, p<0.05, respectively; Figures 5A, 6A and 6B). Regarding the leptin receptor, the IHC score was also significantly higher in animals with a luminal B tumour subtype than healthy animals (2.50 vs. 1.75; p=0.0425; Figures 5B, 6C and 6D).

**Figure 5.**
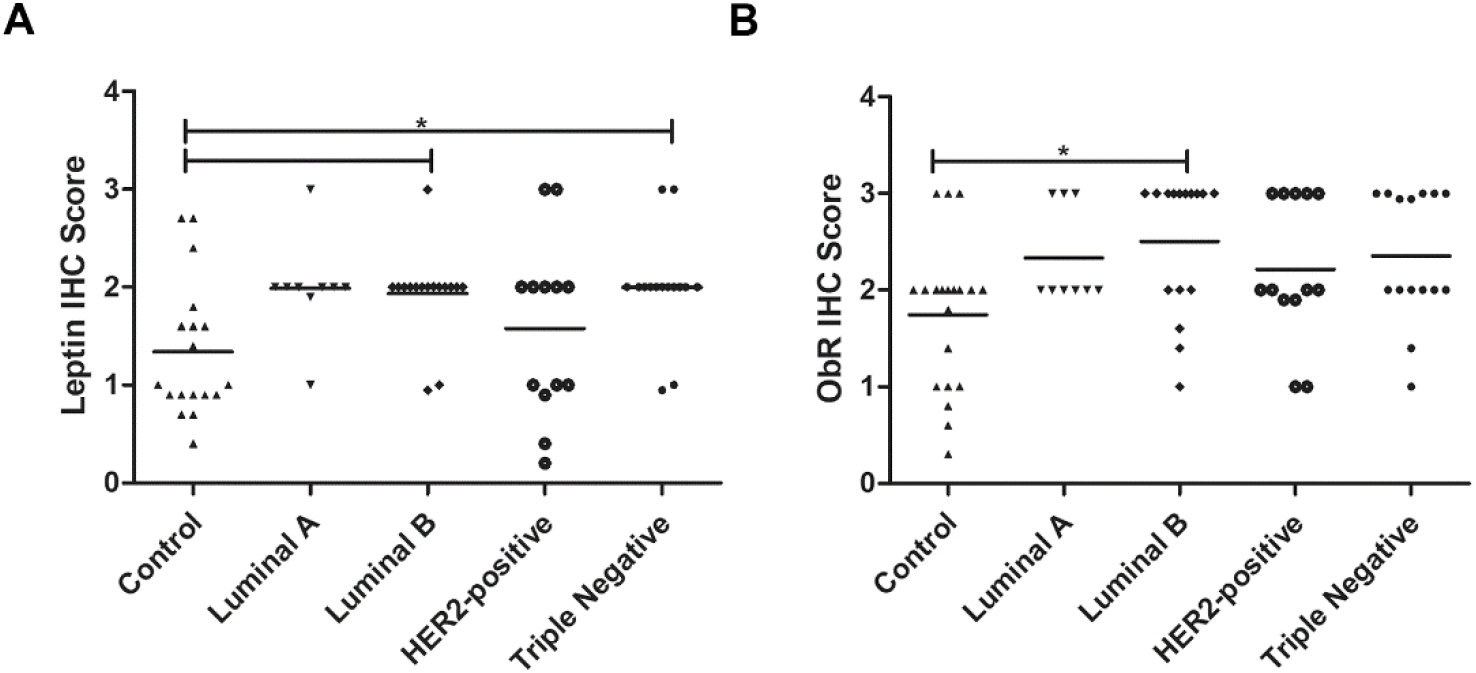
Final IHC scores for leptin **(A)** and ObR **(B)** in cats with mammary carcinoma stratified by tumour subtype and controls. **A)** Leptin expression was significantly higher in luminal B and triple-negative subtypes (p=0.0046). **B)** Expression of ObR was statistically higher in luminal B tumour subtype (p=0.0425).

**Figure 6.**
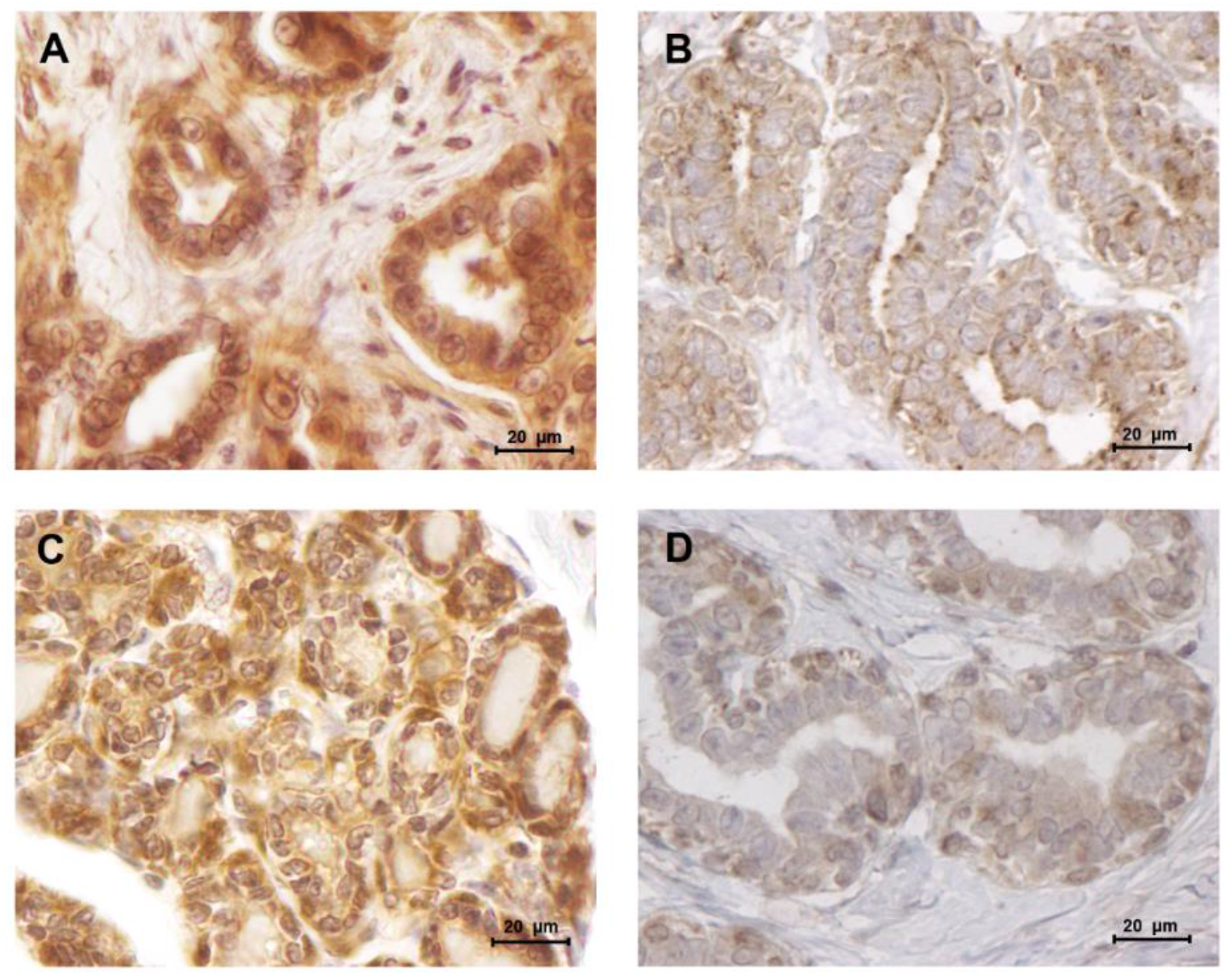
Leptin and ObR are overexpressed in luminal B mammary carcinomas. **A)** Leptin overexpression in a luminal B mammary carcinoma (IHC score of 1.93) contrasting with **B)** a low staining intensity detected in normal mammary tissues (IHC score of 1.34). **C)** Luminal B mammary tumours showed a higher staining intensity for ObR (IHC score of 2.50), **D)** than normal mammary tissues (IHC score of 1.75). (400x magnification)

In addition, our findings revealed that serum ObR levels are negatively correlated with the ObR IHC score, with cats presenting higher serum ObR levels showing mammary tumours with lower ObR IHC scores (p=0.0103, Figure 7).

**Figure 7.**
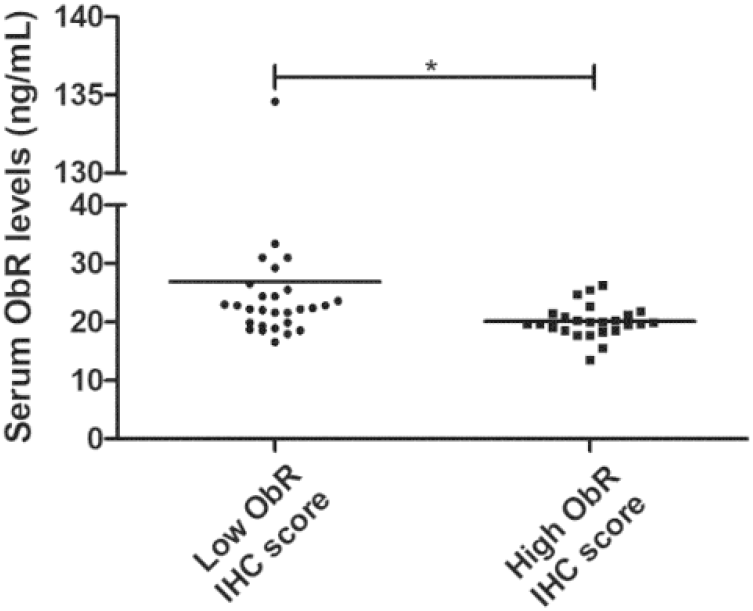
Dot plot diagram showing a negative correlation between serum ObR levels and tumour ObR IHC score (p= 0.0103).

## 4. Discussion

Although spontaneous feline mammary carcinoma has been proposed as a suitable model for the study of human breast cancer, the role of the leptin/ObR axis has never been exploited in cats with mammary carcinoma. In humans, previous studies showed that leptin and ObR overexpression are associated with pro-inflammatory and pro-tumorigenic effects, particularly in overweight women [7] [5].

The results obtained in this study showed that cats with mammary carcinoma have a reduced Free Leptin Index (FLI) in comparison to healthy animals (p=0.0006), suggesting that diseased animals may have decreased soluble leptin levels, as reported in pre-menopausal women with breast cancer [37], and colon cancer patients [38], indicating that serum leptin may be recruited by mammary cancer cells to promote tumour growth and cell migration [11]. Indeed, cats with luminal B or HER2-positive mammary carcinoma showed significant lower serum leptin levels when compared with controls (p<0.001 and p<0.05), revealing that serum leptin levels are downregulated in tumours with PR-positive status [6] and/or HER2-positive status [14]. In contrast, cats with luminal A showed elevated serum leptin levels, suggesting that ER overexpression in the tumour may promotes leptin expression [6]. Regarding the elevated serum leptin levels found in cats with triple-negative mammary carcinomas, studies demonstrated that leptin induces cell proliferative capacity (e.g. via Wnt/β-catenin pathway) [30] and promotes cell survival by interacting with Bcl-2 proteins, being associated with more aggressive tumours [39]. Indeed, elevated serum leptin levels were significantly associated with tumour ulceration (p=0.0005) and shorter DFS (p=0.0217), as reported for breast cancer patients [6] [7].

In parallel, serum ObR levels can also affect the FLI [40], and as reported in breast cancer patients [6] [41], all cats with mammary carcinoma showed higher serum ObR levels than healthy controls (p<0.0001), being correlated with smaller tumour size (p=0.0118) and suggesting that ObR shedding occurs in small tumours, modulating the serum levels of free leptin [15]. Indeed, our results further support the hypothesis that malignant cells in larger tumours maintain the ObR expression on its surface to increase their survival and growth [8]. Interestingly, the higher serum ObR levels were found in cats with mammary carcinomas showing both HER2-positive and ER-negative status (p=0.0291), as reported for human breast cancer patients [6], confirming the crosstalk between the leptin/ObR axis and the EGFR downstream signalling pathway [42].

In addition, this study discloses the utility of leptin and ObR as promising diagnostic biomarkers for feline mammary carcinoma, with a cut-off value of 4.17 pg/mL determined for serum leptin levels to predict feline mammary carcinoma (AUC=0.7045±0.0757, sensitivity=96.9%, specificity=43.5%), whereas a cut-off value of 16.89 ng/mL was calculated for serum ObR levels (AUC=0.9408±0.0288, sensitivity=94.8%, specificity=87.0%) to differentiate animals with FMC from healthy cats.

Interestingly, we also found that serum ObR levels were positively correlated with serum CTLA-4 (p=0.0056), TNF-α (p=0.0025), PD-1 (p=0.0023) levels as reported in breast cancer [26], and with serum PD-L1 levels (p=0.0002). Indeed, previous studies showed that activation of the leptin/ObR axis can result in a chronic inflammatory status [22] [43], a well-known risk factor for breast cancer, with leptin being involved in CD4+ T-regulatory cells differentiation due to ObR overexpression on lymphocyte plasm membrane [44]. These activated CD4+ T-regulatory cells express CTLA-4 [22] and PD-1, two immune-inhibitory checkpoint molecules that downregulate T-cell immune responses [24], leading to tumoral development [45] and contributing to cell growth [46]. On the other hand, in an attempt to control the tumorigenesis process, CD4+ T-regulatory cells secrete TNF-α [25], a molecule that shows a dual role in immunomodulation, being also expressed by cancer cells [47] acting as an autocrine growth factor [48]. Altogether, these findings provide support to the crosstalk between the leptin/ObR axis and tumour immunoediting mechanisms, contributing to an immunosuppressive status in cats with mammary carcinoma [35].

The immunostaining analysis of the tumour and normal tissue samples revealed that luminal B and triple-negative mammary carcinoma subtypes (p<0.05) showed leptin overexpression, whereas a strong ObR expression was detected in luminal B mammary carcinomas (p=0.0425), as described in human breast cancer [13], with several studies suggesting that leptin and ObR are overexpressed in tumoral tissues, due to hypoxia and/or as a response to insulin, IgF-1 and/or to oestradiol [41]. In addition, the higher IHC scores for leptin found in luminal B carcinomas also support the previously reported association between the expression of this adipocytokine and aromatase expression, an enzyme that catalyses the conversion of androgen into oestrogen to promote tumour development via an ER-dependent mechanism [6]. The overexpression of leptin detected in triple-negative mammary carcinomas is also in concordance with previous results in triple-negative breast cancer, where leptin signalling is crucial for tumour growth [16] [39], being associated with ERK and Akt pathways, both involved in breast cancer cells proliferation [11]. Finally, our results demonstrated that cats with low ObR-expressing mammary tumours had higher serum ObR levels, indicating a negative feedback between tumour microenvironment and serum, probably due to a shedding mechanism that leads to a reduction of serum leptin levels [11] [40]. Furthermore, the data obtained emphasizes the possibility to block the leptin/leptin receptor axis, as an adjuvant therapy in cats with luminal B and triple-negative tumour mammary carcinoma subtypes, as reported for breast cancer patients [29] [32] [31].

## 5. Conclusion

This study evaluated the serum leptin/ObR levels in cats with mammary carcinomas providing rationale for their use as diagnostic biomarkers and, additionally, the utility of serum leptin levels as prognostic indicator for the disease-free survival. Indeed, cats with mammary carcinoma showed a significant reduction in the FLI, coupled with decreased serum leptin levels in cats that had luminal B or triple-negative mammary carcinoma and significant increased serum ObR levels, independently of the tumour subtype. As discussed, serum leptin levels above the cut-off value of 4.17 pg/mL were associated to a shorter DFS, whereas serum ObR levels above 16.89 ng/mL were associated to an immunosuppressive status. In tumour tissue samples, leptin is highly expressed in luminal B and triple-negative mammary carcinomas, with ObR being overexpressed in luminal B subtype. Altogether, the data presented extend the knowledge about the similarities between FMC and human breast cancer, further supporting the utility of spontaneous feline mammary carcinoma as a model for comparative oncology studies.

## Author Contributions

Conceptualization, A.G. and F.F.; Methodology, A.G. and F.F.; Formal analysis, A.G., C.N.; Investigation, A.G., C.N., A.C.U., J.C and F.F; Supervision, F.F.; Funding acquisition, F.F; Project administration, F.F.; Writing – original draft preparation, A.G. and F.F.; Writing – review and editing, A.G., C.N., A.C.U., J.C., and F.F. All authors have read and agreed to the published version of the manuscript.

## Funding

This research was funded by Fundação para a Ciência e a Tecnologia (Portugal) through the projects PTDC/CVT-EPI/3638/2014 and CIISA-UID/CVT/00276/2020. A.G. is receipt of a PhD fellowship from Fundação para a Ciência e a Tecnologia (ref. SFRH/BD/132260/2017) and C.N. is receipt of a PhD fellowship from University of Lisbon (ref.C00191r).

## Acknowledgments

The authors would like to thank Dra. Maria Soares for the clinical samples and database.

## Conflicts of Interest

The authors declare no conflict of interest. The funders had no role in the design of the study; in the collection, analyses, or interpretation of data; in the writing of the manuscript, or in the decision to publish the results.

## Notes

### Competing Interest Statement

The authors have declared no competing interest.

